# MVD-Fuse: Detection of White Matter Degeneration via Multi-View Learning of Diffusion Microstructure

**DOI:** 10.1101/2021.04.15.440095

**Authors:** Shreyas Fadnavis, Pablo Polosecki, Eleftherios Garyfallidis, Eduardo Castro, Guillermo Cecchi

**Affiliations:** IBM T.J. Watson Research Center; Indiana Univeristy Bloomington

## Abstract

Detecting neuro-degenerative disorders in early-stage and asymptomatic patients is challenging. Diffusion MRI (dMRI) has shown great success in generating biomarkers for cellular organization at the microscale level using complex biophysical models, but there has never been a consensus on a clinically usable standard model. Here, we propose a new framework (MVD-Fuse) to integrate measures of diverse diffusion models to detect alterations of white matter microstructure. The spatial maps generated by each measure are considered as a different diffusion representation (view), the fusion of these views being used to detect differences between clinically distinct groups. We investigate three different strategies for performing intermediate fusion: neural networks (NN), multiple kernel learning (MKL) and multi-view boosting (MVB). As a proof of concept, we applied MVD-Fuse to a classification of premanifest Huntington’s disease (pre-HD) individuals and healthy controls in the TRACK-ON cohort. Our results indicate that the MVD-Fuse boosts predictive power, especially with MKL (0.90 AUC vs 0.85 with the best single diffusion measure). Overall, our results suggest that an improved characterization of pathological brain microstructure can be obtained by combining various measures from multiple diffusion models.

## 1 Introduction

Diffusion-weighted magnetic resonance imaging (dMRI) exploits the diffusion of water along white matter pathways in the brain to quantify its microstructure and reconstruct its major tracts in the living human brain [4]. To probe microstructure using DWI, the signal representations are typically obtained by decomposing the 4-dimensional signal (at a voxel scale) using tensor, multi-compartmental or spherical harmonic models. Measures derived from modeling DWI typically reveal information related to degree of coherence of neuronal fiber alignment, microstructural integrity, demyelination and corruption of microtubules.

A wide range of models have been proposed to probe the tissue microstructure, yet, there has been no consensus on a particular standard model and no single model can provide an accurate description of the underlying signal structure to detect pathological changes of white matter [9, 12]. Each microstructure model provides unique information about the tissue structure describing the same tissue degeneracy. This represents our primary motivation for fusing information derived from multiple models: to learn measures specific to particular diseases. In this work, we investigate multi-view machine learning methods to integrate the predictive information from all DWI models in the diagnosis and stratification of neurological disorders.

We propose a new framework, Multi-View learning of Diffusion Microstructure (MVD-Fuse), which contains the following innovations: (1) MVD-Fuse aims at characterizing white matter alterations by optimizing for predictive accuracy for specific disorders, as opposed to the massive univariate analyses that historically dominated the field [13]. (2) The proposed framework fuses measures from all models exploiting state-of-the-art multi-view fusion approaches. In terms of the ability to learn optimal fused representations, we compared the performance of algorithms from two broad categories: early and intermediate fusion (EF and IF, respectively). EF consists of concatenating all views into a single feature vector for training, while IF implies learning linear combinations 34th Conference on Neural Information Processing Systems (NeurIPS 2020), Vancouver, Canada. of the input views in the latent space, which better captures the interaction between individual views. Additionally, IF allows the ranking of model contributions by inspecting the view-specific weights of the learnt representations. We explored three fundamental approaches to IF: Multiple Kernel Learning (MKL) [7], Muli-View Boosting (MVB) [8] and Neural Networks (NN) [3].

In this paper, we work with Huntington’s disease (HD) as a case study. HD is a monogenic disease which can be detected via genetic tests, enabling large observational studies on the HD-gene carrying subjects before diagnosis based on symptom manifestation [11]. Specifically, we evaluate MVD-Fuse’s ability to detect differences in white matter microstructure between premanifest HD individuals (pre-HD) and healthy controls.

## 2 Data and Methods

### Data

We evaluate the ability of MVD-Fuse to distinguish pre-HD subjects from controls in the TrackON dataset [10]. The data in this work contained 140 subjects (68 healthy controls, 72 pre-HD). dMRI data was acquired using a single-shell acquisition protocol with 41 gradient directions at b-value 1000 each. The data came from four different imaging sites. Therefore we followed a cross-site cross-validation approach to ensure robust generalization across sites [1].

### Methods

The proposed framework starts by fitting the dMRI data with different microstructure models. Here, we fit the data with three widely used microstructure models, namely, constrained spherical deconvolution (CSD) [16], diffusion tensor (DTI) [4] and the single-shell free-water diffusion tensor models (FW-DTI) [14]. FW-DTI is a two-compartmental model that provides information about signal contributions from extracellular free water (FW). DTI on the other hand is a single compartment model aimed at deriving information about intrinsic mean diffusivity (MD) and fractional anisotropy (FA) of the diffusion process. As opposed to phenomenological models like DTI and FW-DTI, CSD takes a mechanistic approach to deconvolve the signal and derive fiber Orientation Distribution Function (fODF) using spherical harmonic expansions, giving measures such as Generalized FA (GFA) and Apparent Fiber Density (AFD) [see Fig. 2A].

**Figure 1:**
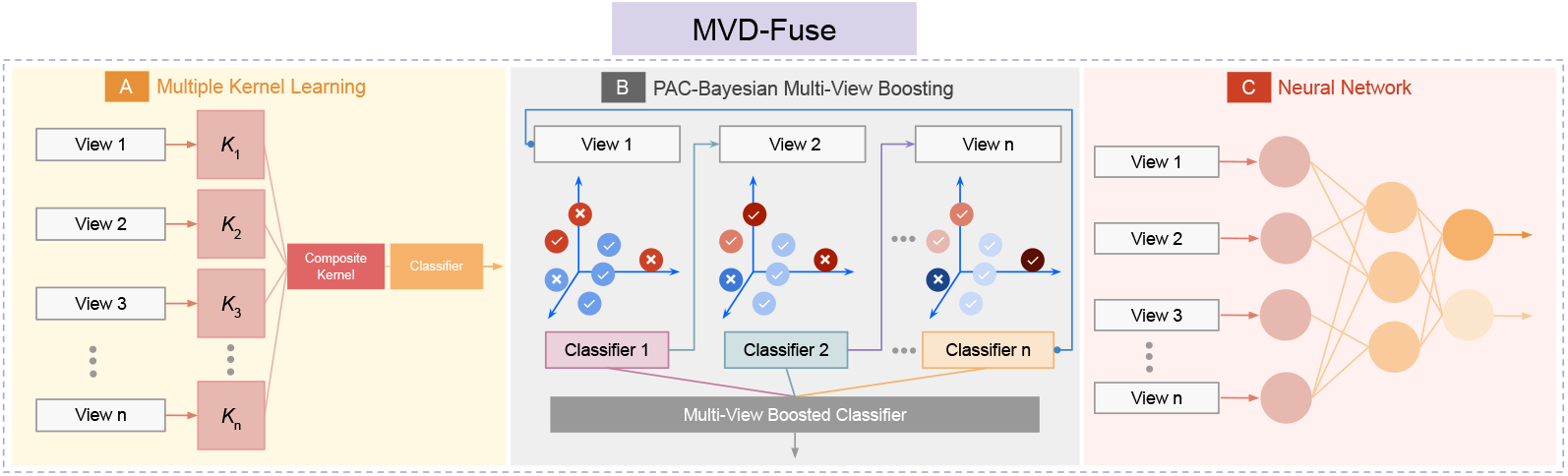
MVD-Fuse framework. (A) MKL, (B) MVB and (C) NN intermediate fusion algorithms are used to integrate different measures (views) derived from DWI modeling and improve the detection of pathological patterns of white matter microstructure in a disorder of interest (pre-HD in our proof-of-concept validation).

**Figure 2:**
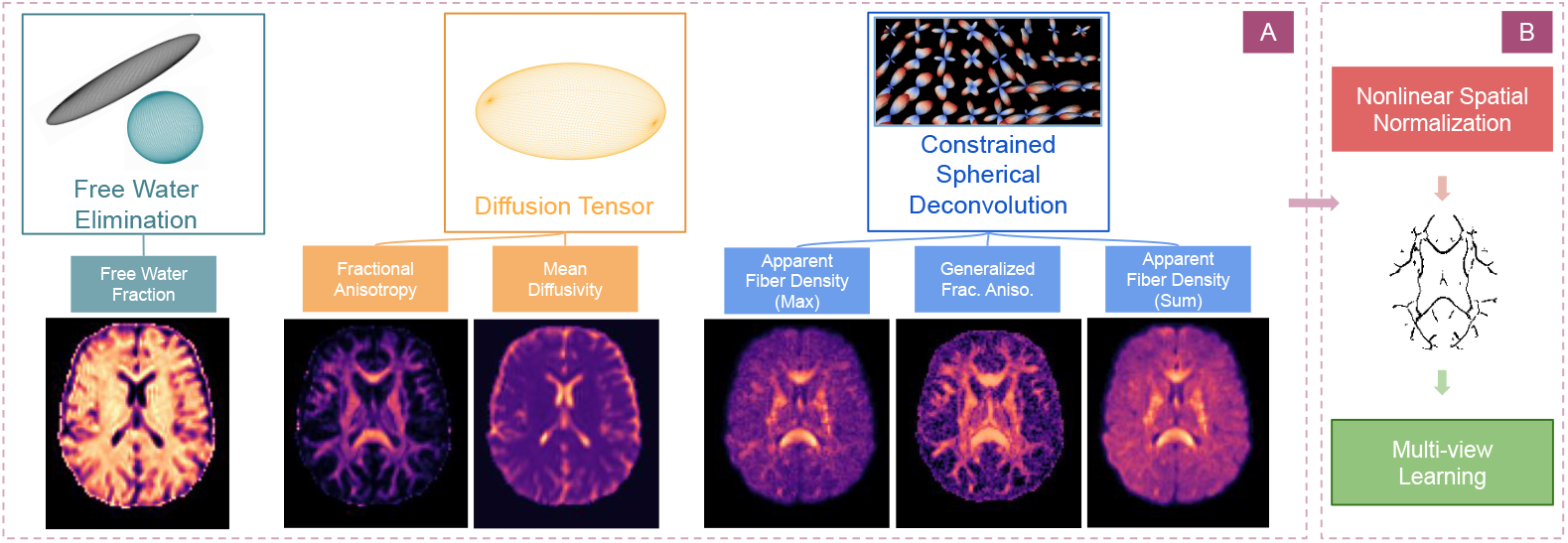
Depiction of analyzed measures and extracted features. **(A)** Models fitted to the data and microstructure maps derived as outputs from them. **(B)** Nonlinear registration of each model fit to a white matter template, which is in turn fed to the multi-view learning algorithm as input.

After model fitting, subjects are registered to a standard WM FA template [15]. Then, a common WM skeleton is computed (tract-based spatial statistics, TBSS) and all measures are projected onto it. We additionally removed skeleton voxels where FA had an inter-subject coefficient of variation above 0.3, retaining a core with minimal anatomical heterogeneity. Measures from all models along the WM skeleton are then fed into the multi-view learning framework to perform the binary classification task of distinguishing pre-HD subjects from healthy controls. When using an EF approach, all the derived measures are concatenated into a single feature vector, discarding information about the diffusion model from which each feature originates. We applied standard linear classifiers to train the concatenated features, mainly Support Vector Machines (SVM) and Elastic Net Logistic Regression.

As a first IF approach, we explored a dense 2-layer linear NN, where features from each view feed a view-specific unit in an intermediate layer. These intermediate layers learn a view-specific representation, which is fused by an output unit. In MKL, each of the microstructure measures were input to linear kernels (*K*_1_, *K*_2_, … *K_n_*) as shown in Fig. 1A. Then, all the kernels corresponding to the different views were fused together using the EasyMKL [2] algorithm to learn the optimal linear combination of these kernels. Next, we used the SVM classifier [5] to perform the classification using the resulting composite kernel. Lastly, we explored MVB [8], which is an extension of Adaboost [6] for multiview integration. MVB uses a hierarchical strategy: it first estimates weights for each of the base weak-learners (here decision trees) generated within a given view, and then estimates view-specific weights. This is done by controlling a trade-off between classification error and prediction diversity among views. We use the receiver operator characteristic (ROC) curves along with area under the ROC curve (AUC) scores to assess the performance of these classification approaches.

## 3 Experiments and Results

First, we evaluated the predictive power of single measures as a baseline for comparing against multi-view models (Fig. 3A, Table 1). Each individual measure had significant discriminative ability. Next, we used MVD-Fuse with different multi-view methods (IF) and compared them against EF-based classifiers. For EF, elastic net logistic regression gave an AUC score of 0.77 and the support vector classifier gave an AUC score of 0.83. NN and MVB did not outperform the SVM (EF) classifier but. On the other hand, MKL improved the predictive accuracy by 5% points (AUC score of 0.90), showing better performance than any single model and EF approach.

**Figure 3:**
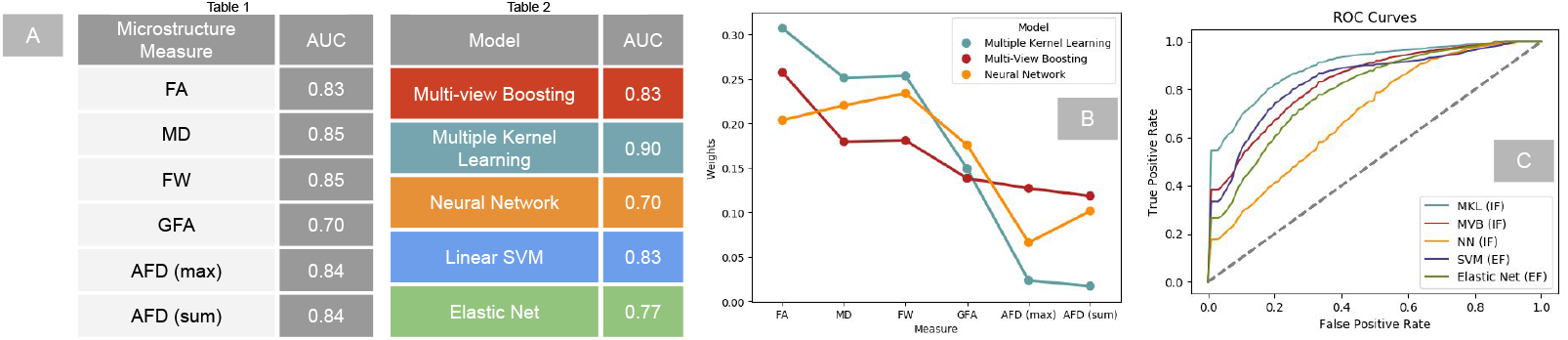
Overall performance of the models and assessment of informative dMRI measures. (A) AUC performance of single measures (Table 1) and the different strategies for multi-view learning (Table 2). (B) View-specific contributions learnt from each of the multi-view methods in MVD-Fuse. (C) ROC curves of different methods.

## 4 Conclusions

In this work, we propose a novel way of integrating microstructure information that differs from traditional model selection, in a way akin to how multivariate machine learning differs from univariate tests. This not only avoids the need for separate rounds of model selection and prediction, which are prone to “double dipping”, but enables integration of complementary information, thus boosting predictive power. Moreover, our results are interpretable, as shown by view-specific weights that were consistent across IF approaches. Detailed future work will investigate how this multiview integration can produce composite voxel-level measures optimally sensitive to alterations from specific disorders.

## 5 Broader Impacts

This work introduces information fusion via multi-view learning enabling disease detection and potentially can be used for predicting progressions. This approach can naturally be extended to incorporate other modalities such as Perfusion MRI, Functional MRI, MEG/ EEG, etc. Relatively, there is lack of methods to perform multi-modal analyses and this work sheds light upon its usefulness from an application standpoint. Beyond the medical imaging community, MVD-Fuse implements the general idea of decomposing the same data to generate embeddings with unique information that can be used as features for any downstream machine learning task.

